# Metabolic Implications of Birth via Cesarean Section in Prairie Vole (*Microtus ochrogaster*)

**DOI:** 10.1101/2022.05.21.492929

**Authors:** Alexandra Starr, Sabreen Ahmed, Miranda Partie, William Kenkel

## Abstract

Throughout the United States the rates of performed cesarean section (CS) have increased. The scientific community has observed an association between birth by cesarean and the offspring’s increased weight at maturity (Masukume, 2019). Studies are being conducted to better understand the relationship between cesarean delivery and offspring metabolism (Kozhimannil, 2013; Kenkel, 2020). To test this potential connection, a diet intervention study has been used to test vaginal delivery (VD) vs CS birth subject’s weight gain using a prairie vole model. Vole diets were either supplemented with a high-fat alternative mixed chow (MC) or fed standard vole chow (VC) to induce weight gain. Through this study, we collected sucrose preference, home cage, food consumption data from both birth mode groups and diet conditions. At sacrifice, we collected measures of weight, length, and adipose tissue to analyze for post-mortem body composition in adulthood of each group. CS voles gained more weight than VD voles, despite having lower food consumption and greater locomotive activity. Body composition analysis found that CS animals were longer and heavier than their VD counterparts. Additionally, CS animals were found to have a larger percent brown adipose tissue relative to body weight compared to VD counterparts. Future studies will target the variables contributing to this weight gain among CS offspring by examining factors like muscle mass, and total adiposity through advanced imaging data. Future studies will incorporate exogenous oxytocin administration to examine the impact of birth mode on body weight, metabolism, adiposity, and later life development to determine the possible mechanisms impacting the metabolic outcomes seen in this study.

## INTRODUCTION

### 1.1 Cesarean Section

#### 1.1.1 History of Cesarean Section Use and Procedures

Records of birth via cesarean section (CS) have been found in literature from many ancient societies across history. Cesarean Section was originally introduced as a procedure to retrieve the infant from a dead or dying mother post-mortem. The frequency of Cesarean Section increased with political and religious influence with the logic that a pregnant woman was forbidden burial until the child had been removed– and the two blessed and buried separately (NIH, 2013). The term cesarean section has often been linked to the birth of Julius Caesar in ancient Rome which is apocryphal as Julius Caesar’s mother is recorded to have survived childbirth dying many years later (Fadel et al., 2011). It is more likely that the term arose from the Roman legal code depicted in the Lex Regia passed in 715 BCE Lex Regis de lnferendo Mortus. The law forbade the “burial of a pregnant woman until the child had been removed from her abdomen, even when there was but little chance of its survival, so that the child and mother could be buried separately.” (Fadel et al., 2011). Regardless of the Cesarean section’s nomenclature or historical origins the procedure increased in frequency and precision with the medical developments of the 16th century and the mass printing of medical texts across Europe. With the medical advancements of the 19th and 20th century allowed for the improved survival of both the mother and child and by the beginning of the 20th century cesarean sections began to resemble what we would find in today’s operating suites.

#### 1.1.2 Modern Cesarean Section

Until the 20th century most births occurred in the home under the care of midwives and family; doctors were only involved in the case of emergency (Scott, 2011). By the 1950s 88% percent of births occurred in a hospital with a physician instead of at home with a midwife – this number continued to increase until 2017 where approximately 98.4% of births were done in hospital (MacDorman, 2011). Initially introduced as a life-saving practice cesarean procedures are now being used preemptively in the case of a challenging labor to save both mother and child (Jansen, 2013).

As of 2015 cesarean sections (CS) account for one third of deliveries across the United States (Montoya-Williams, 2017). Internationally, a 2010 WHO report of CS rates in 137 countries found a trend that within countries in Africa and Asia CS rates were approximately less than 10% and in countries like Brazil and Iran CS births represented nearly half of all births (World Health Organization Human Reproduction Programme, 2015). While CS procedures are usually medically necessary and frequently lifesaving, C-sections are considered a major surgical procedure and are associated with many health risks to mother and fetus, including cardiac arrest, hysterectomy for the mother, and maternal and fetal mortality (Liu, 2007). There is increasing evidence that children born via CS experience higher rates of adverse health outcomes later in life (Yuan, 2016). One of the most well-established associations between CS delivery and offspring development is increased weight later in life (Masukume, 2019). In 2008, a North American study reported that children born by CS are 40% more likely to be overweight (Sogunle, 2019). Obesity is a condition characterized by excessive body fat that increases the risk of developing associated health problems. A recent meta-analysis that found CS delivery is associated with a 59% increase in the risk of developing obesity by 5 years of age (Keag et al., 2018). There is evidence of an association between delivery by CS and metabolic outcomes, but it may be driven by undetermined confounding variables. Clarification is necessary to determine any harmful consequences of cesarean delivery on offspring health.

### 1.2 Rodent Model

Human clinical research is important but cannot often account for the many limitations and variables due to the complex and diverse lifestyle of human subjects. Human research in c-sections is not able to control for conditions such as maternal obesity, overall parental health and gestational age (Masukume, 2018). However animal models allow for total control during the study and operate on a more expedited timeline. The prairie vole model is a better choice than the traditional mice model in studying metabolism. Prairie voles are better suited to room temperature housing, cross fostering, and show an equivalence between the sexes. Rodents in general and prairie voles in particular are very energy efficient and are unlikely to develop obesity in a lab setting, thus it is necessary to administer a high fat diet to induce weight gain (Martin et al., 2010). The current study of cesarean section and its impact on metabolic outcomes consists of two facets of interest: high fat diet induced weight gain and the accumulation of visceral lipids.

In studies of metabolism, it is critically important to consider thermoregulation as an important factor in small mammal energy budgets. In the past, this has stymied rodent studies of diet-induced obesity (Cui et al., 2016; Feldmann et al., 2009; Giles et al., 2017; Stemmer et al., 2015)because traditional laboratory rodents experience conventional 20°-22° housing as a chronic cold stress (Ganeshan & Chawla, 2017; Kokolus et al., 2013; Maloney et al., 2014a). Housing temperature is one the largest determinants of energy expenditure in mice (Corrigan et al., 2020). Indeed, basal metabolic rate and food intake of mice housed at 20° are both ~50% higher than those housed at a thermoneutral 30° (Cui et al., 2016; Giles et al., 2017; Hankenson et al., 2018; Maloney et al., 2014b). Thus, the goal of our first experiment in this study was to investigate the impact of ambient temperature of prairie vole metabolism. We sought to investigate whether prairie voles are as sensitive to ambient temperature as traditional lab rodents.

#### 1.2.1 Body Composition

The assessment of body fat and associated body composition is essential for the diagnosis of obesity (Hu et al., 2012). While it is challenging to make a claim of obesity in an animal model, it is easier to evaluate animals for the most characteristic factor of obesity, which is weight gain. Without a complex imaging technique, it is possible to use simple measures to indirectly measure weight and body composition in the evaluation of health in a rodent model. One method by which to calculate health is the Body Mass Index (BMI). The BMI calculation involves measures of weight and length to assess health relating to obesity. Recent studies have found a positive correlation between BMI and carcass fat in rats (Novelli et al.,2007). This idea demonstrates a possible link between BMI and the measure of obesity and allows us to indirectly measure the health and body fat of a rodent model. The Lee’s index is another calculation used in animal models to assess body composition. The Lee’s index is an older method of calculation that does not correlate well with other standard measures of body composition (Rogers et al., 1980). Along both of these indirect calculations of body fat, the proportional weight of adipose tissue samples or ‘fat pads’ is widely recommended as a simple and direct estimate of body fat in normal or high-fat diet obesity induced rodent models.

As depicted in Figure 1, smaller mammal species have two major types of adipose tissue: brown adipose (BAT) and white adipose tissue (WAT)(Almind et al., 2007). The white adipose tissue (WAT) is composed of two subcutaneous pads (anterior and posterior). WAT is more widely distributed throughout the body and stores excess energy as triglycerides. BAT is physiologically important due to its importance in metabolic regulation (Townsend et al., 2012). Not pictured in the figure are the several smaller intramuscular and retroperitoneal fat deposits that develop over time. These two fat pads have been used as a representational sample to correlate to total adiposity of the animal in rodent models. A 2016 study examining the *Body fat of rats of different age groups and nutritional states: assessment by micro-CT and skinfold thickness* found a significant relationship between certain wet weights of adipose tissue samples and abdominal micro computed tomography (micro-CT) derived total adipose tissue mass in a rodent model (Takus et al., 2016). This study and others demonstrated how the weight of certain adipose tissue samples can be used to assess whole body composition with relative accuracy.

**Figure 1:**
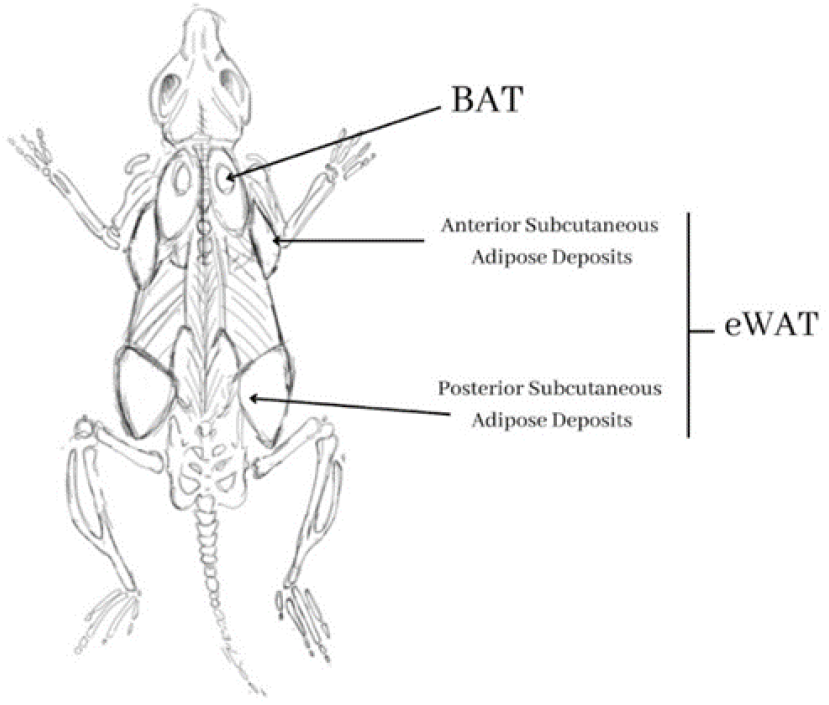
Distribution of Fat pads in Prairie Vole: The anterior and posterior subcutaneous WAT pads are located between the scapulae and the dorsolumbar-gluteal region respectively. The BAT is distributed throughout the viscera with a localized fat pad in the interscapular region depicted to the left.

### 1.3 The Diet Intervention

There are several methods to induce weight gain in a rodent model. For this study a diet-induced obesity procedure was used to investigate the Animal’s weight gain in VD and CS offspring. In studies of both mice and rats, a positive correlation has been found between the level of fat in the diet and a voles’ fat content or body weight. To examine the obesogenic effect of CS on prairie vole, it is important to determine the effect of supplementing the voles’ diet with a High-Fat mouse chow alternative (MC) with the intention to induce weight gain.

### 1.4 Sucrose Preferences

Sucrose preference is associated traditionally with a test of appetite regulation. In this study the test of sucrose preference at weaning (baseline) and at adulthood examines the impact of CS offspring birth on preferential consumption of sucrose solution. Consumption behavior is an important variable when examining the differences between birth mode groups and diet groups. In rodent models, responsiveness to sucrose, but not fat, was an effective predictor of weight and adiposity gain (Grinker et al., 1978). In a study from 2006, research in humans found that subjects contending with generalized obesity display a greater preference or ‘liking’ for sweet solutions compared to non-obese subjects (Bartoshuk et al, 2006). Numerous studies have not found a definitive role in sucrose preference to the susceptibility to obesity and the high fat diet.

### 1.5 Rationale for Current Research

In this study we examined the impact of birth mode on body weight, metabolism, adiposity, and later life development of the prairie vole (*Microtus ochrogaster).* This study compared prairie voles delivered by cesarean section (CS) to those delivered vaginally (VD) throughout development into adulthood. We tested these birth mode groups against two diet conditions; a standard chow (VC) and a mixed high-fat diet (MC). To study the impact of birth mode on metabolism, we tested several behavioral, nutritional, and physiological measures on both diet groups (MC and VC).

The period of testing included measures of food consumption, sucrose preference, and home cage activity. Among these tests, we predicted that CS animals would have increased sucrose preference compared to their VD counterparts, due to the association between sucrose preference levels and appetite. We hypothesized that CS offspring would gain more weight than their VD counterparts and additionally consume greater quantities of food across both diet conditions. Our hypothesis was rationalized by the finding that CS delivery is associated with a 59% increase in the risk of developing obesity by 5 years of age (Keag et al., 2018). We also predicted this weight gain would be associated with greater food intake by the CS group and decreased locomotor activity in the home cage. This total body weight gain was further evaluated by several factors–including: total weight, length, mass of representational tissue samples, BMI and Lee’s Index. We measured length to control for differences in body weight resulting from increased animal length. We hypothesized that CS delivery would be associated with higher BMI and higher Lee’s Index score. Associated with this hypothesis, we predicted to find greater mass of adipose tissue samples, proportional to the animals weight, in CS offspring in both BAT and WAT samples.

## 2 METHODS

This study consisted of two experiments. In the first, we sought to examine prairie voles’ metabolic responses to varying ambient temperature. In the second, we sought to examine prairie voles’ metabolic responses to being delivered via cesarean section.

### 2.1 Experiment 1

In Experiment 1, 7 adult prairie voles (3 female and 4 male) housed at the University of Delaware were used for this study. Subjects were subcutaneously implanted with radiotelemetry transmitters (C19BTA, Indus Instruments Inc.). These devices weigh 2.7 grams, measure 1.9 cc in volume, and are compatible with rodents as small as 25 grams (adult voles typically weigh >30 grams). Surgical methods for the implantation of these devices followed those of our previous work (Kenkel et al., 2013, 2014, 2015). Following implantation, subjects were given 10 days to recover in divided cages that prevented both social isolation and interference with healing by cage mates. The following parameters were recorded for 5 minutes every hour: heart rate, locomotor activity, and body temperature.

After the 10 days of recovery, subjects were exposed to different ambient temperature every 72-96 hours using a slightly modified protocol laid out in previous work using rats and mice (Swoap et al., 2008). Testing begin at 22° and incrementally increase by 4° at each time interval to 30°, then incrementally decrease back down to 18°, and finally returning to 30°C. Thus, nine temperatures will used: 22°, 26°, 30°, 26°, 22°, 18°, 22°, 26°, and 30°. During testing, all subjects remained in their home cages throughout the period of temperature fluctuations. Standard nesting and cage mates were maintained throughout the study. Ambient temperature in the vole colony room was maintained and manipulated by adjustments to the thermostat using the Building Automated Systems every 3-4 days so as to avoid changes occurring over weekends. Humidity levels were between 30% and 70%. Subjects were monitored twice daily to ensure they tolerated temperature changes. Finally, subjects’ food and water consumption was recorded daily.

### 2.2 Experiment 2

In Experiment 2, 84 prairie voles (42 female and 42 male) housed at the University of Delaware were used for this study. Subjects were housed by sex (litter sizes ranging from 2-5) in polypropylene cages (dimensions) containing wooden shavings for bedding and various forms of enrichment activities. The housing facility is temperature and light controlled on a 12-hour light/dark cycle, starting from 7am to 7pm.

All measurements and experimentation took place during the light cycle period. Breeding pairs that were used for this study were created in-house. These breeding pairs were not used in any previous experimentation and were evaluated for current level of health, such that they were not noticeably larger or smaller than colony average for approximate age.

21 days after observed mating between the breeding pair, it is necessary to prepare for the birth procedure. When at least one dam within the cohort had been observed completing a spontaneous vaginal birth, another noticeably pregnant dam with no signs of delivery was selected for CS producer. Dams were selected for the CS procedure based on measurable weight gain (approximately 20g) or observable signs of pregnancy. For the CS procedure dams were anesthetized with 2% CO2 and cervical dislocated. Then an abdominal incision was made to expose the uterine horn. The fetuses were removed from the sac and cleaned from their membranes (Castillo-Ruiz et al., 2018). The pups were weighed immediately following birth. Any pups that weighed less than 2.5g were excluded from the study to avoid confounding study with any effects due to preterm birth. Each VD and CS litter with more than 5 pups was culled to 5 during this stage of experimentation. Within 24 hours following vaginal birth, or directly following CS procedure, all offspring were cross fostered to control for possible parental behavioral artifacts.

Postnatal Day 0 (PND0) was designated at the date of spontaneous vaginal birth or procedural c section. Voles were immediately cross-fostered and later weaned at PND21. After weaning, voles were pseudo-randomly assigned to conditions of diet intervention and tested using the following measures: weekly weights, food consumption, sucrose preference and home cage behavioral monitoring. The timepoints of each test conducted are illustrated in Figure 2.

**Figure 2:**
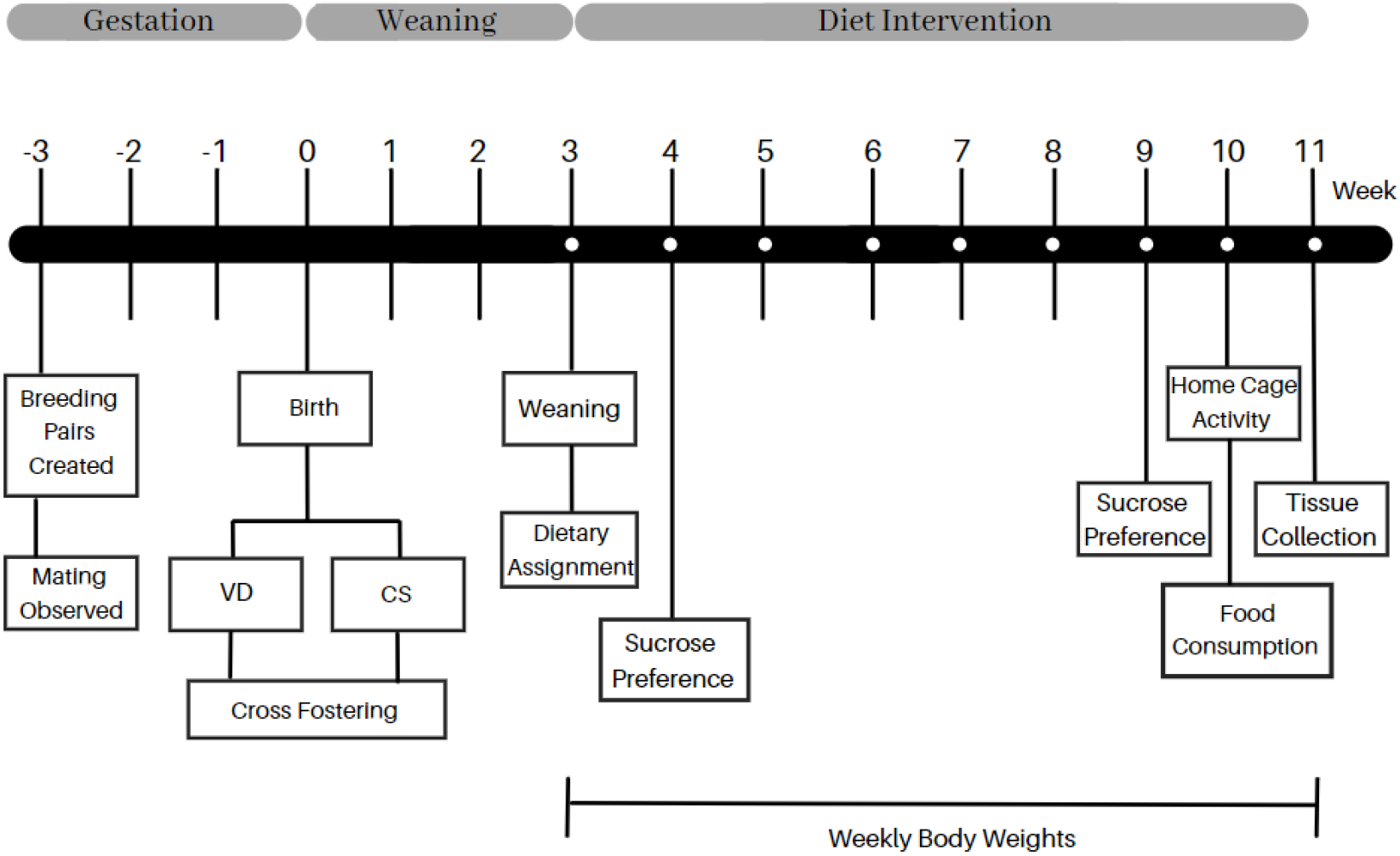
Experimental timeline depicting the series of events and behavioral testing the animals underwent.

During experimentation, food pellets and water bottles were given ad libitum according to the diet intervention assigned at weaning. All practices were approved by the University of Delaware Institutional Animal Care and Use Committee prior to the project’s beginning.

### 2.3 Weekly Weights

All voles were weighed weekly from PND21 to PND75. The scale used was calibrated monthly for accuracy to 0.01grams. Voles were weighed at the same time each day to reduce error introduced through the feeding protocol.

### 2.4 Food - Diet Intervention

At weaning, both CS and VD voles were pseudo-randomly assigned to a diet condition (vole chow or mixed high fat diet chow) based on sex and birth mode. Conditions were divided as evenly as possible by sex of litters in each condition. Vole chow condition (VC) contained 2% vegetable fat, 15% protein, 40–50% carbohydrates, and 15–25% fiber. The mixed high-fat diet (MC) contained approximately 50/50 vole and mouse chow (mouse chow makeup: 7% simple sugars, 3% fat, 50% polysaccharide, 15% protein (w/w), energy 3.5 kcal/g).

#### 2.4.1 Sucrose Preference

Sucrose preference was tested at PND28 and PND63 over a 48-hour period. Animal cages were set up with a “two-bottle choice” procedure consisting of one bottle of 1% sucrose water solution and one bottle of filtered water (Figure 3). Both bottles were weighed at the start of the test and weighed 24-hours later. The position of the 1% sucrose bottle was counterbalanced across birth mode and diet intervention groups to control for location preference effects.

**Figure 3:**
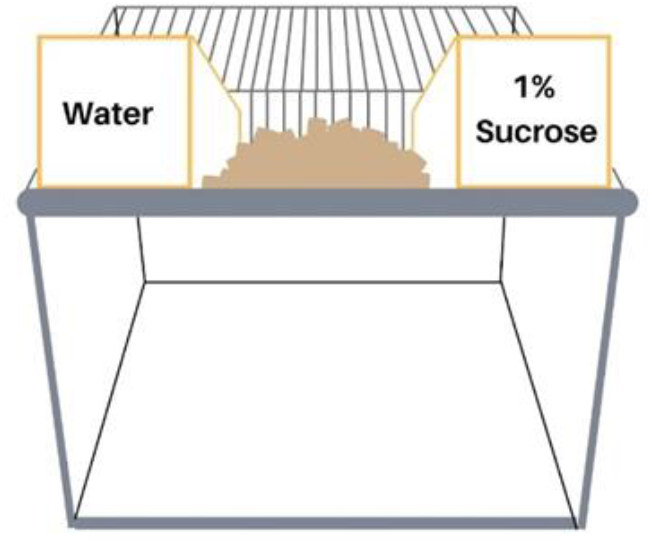
Sucrose Preference Test Cage: with two-bottle preference set up depicting the water sipper bottle on the left and the 1% sucrose and water solution on the right.

#### 2.4.2 24-Hour Representative Food Consumption

At PND70, a 24-hour food consumption test was performed in the home cage for both diet conditions. At 8am, all food pellets were weighed prior to distribution to the home cages. After 24 hours all food remaining in the cage was weighed.

### 2.5 Home Cage Activity Monitoring

At PND71, voles underwent a locomotive observation test for 2 hours in their home cages. Voles were observed by a remote camera suspended above the home cage. Home cage lids were replaced by transparent Plexiglas coverings with air holes to allow for continuous airflow.

### 2.6 Sacrifice and Tissue Collection

At PND75, sacrifice was conducted under routine procedures approved by IACUC using 4% isoflurane anesthesia, followed by cervical dislocation. Vole length and final body weight were recorded. Length of vole was measured from nose to base of tail. After cervical dislocation, necropsy was conducted to collect various tissues for analysis and preservation, including the brain, scapular brown adipose tissue, and anterior and posterior subcutaneous fat pads illustrated in Figure 2. All adipose tissue samples were weighed immediately following excision.

### 2.7 Statistical Analysis

For the food consumption measure, this measure was added to the study in the midst of data collection and had smaller sample sizes; therefore, there were not enough data points to consider sex as a variable. Hence, food consumption was analyzed as a two-way ANOVA with birth mode (CS vs. VD) and diet (MC vs. VC).

For the BAT, WAT, body length, and body weight measures, data were analyzed as a repeated measures linear mixed effect model with litter included as a random variable to account for similarity among siblings.

One VD cage was excluded from analysis due to exceptional weight gain, >2 SDs above the study mean (and greater than any vole of comparable age in colony history). We also excluded 5 VD and 4 CS cages in the MC group as these voles were transferred to a separate pilot experiment.

## RESULTS

### 3.1 Ambient Temperature

We observed a weak relationship between ambient temperature and subjects’ heart rate (Figure 4). Heart rate was negatively correlated with ambient temperature (R = −0.33, p < 0.001). Heart rate was 327 ± 11 beats per minute (bpm) at the warmest temperature (30°), 363 ± 14 bpm at the conventional housing temperature (22°), and 348 ± 16 bpm at the coldest temperature (18°). We observed no order or transition effects, i.e. each instance of a given temperature resembled the other instances of that temperature; it did not matter whether temperature had recently increased or decreased. Ambient temperature was positively correlated with body temperature (R = 0.61, p < 0.001). Body temperature was 36.4 ± 0.1° at the warmest temperature (30°), 35.8 ± 0.1 at the conventional housing temperature (22°), and 35.7 ± 0.2° at the coldest temperature (18°). We observed no effect of ambient temperature on locomotor activity. Finally, we observed no effect of ambient temperature on either food or water consumption, however both measures were carried out on a per cage basis and hence the sample size was reduced (n = 4).

**Figure 4:**
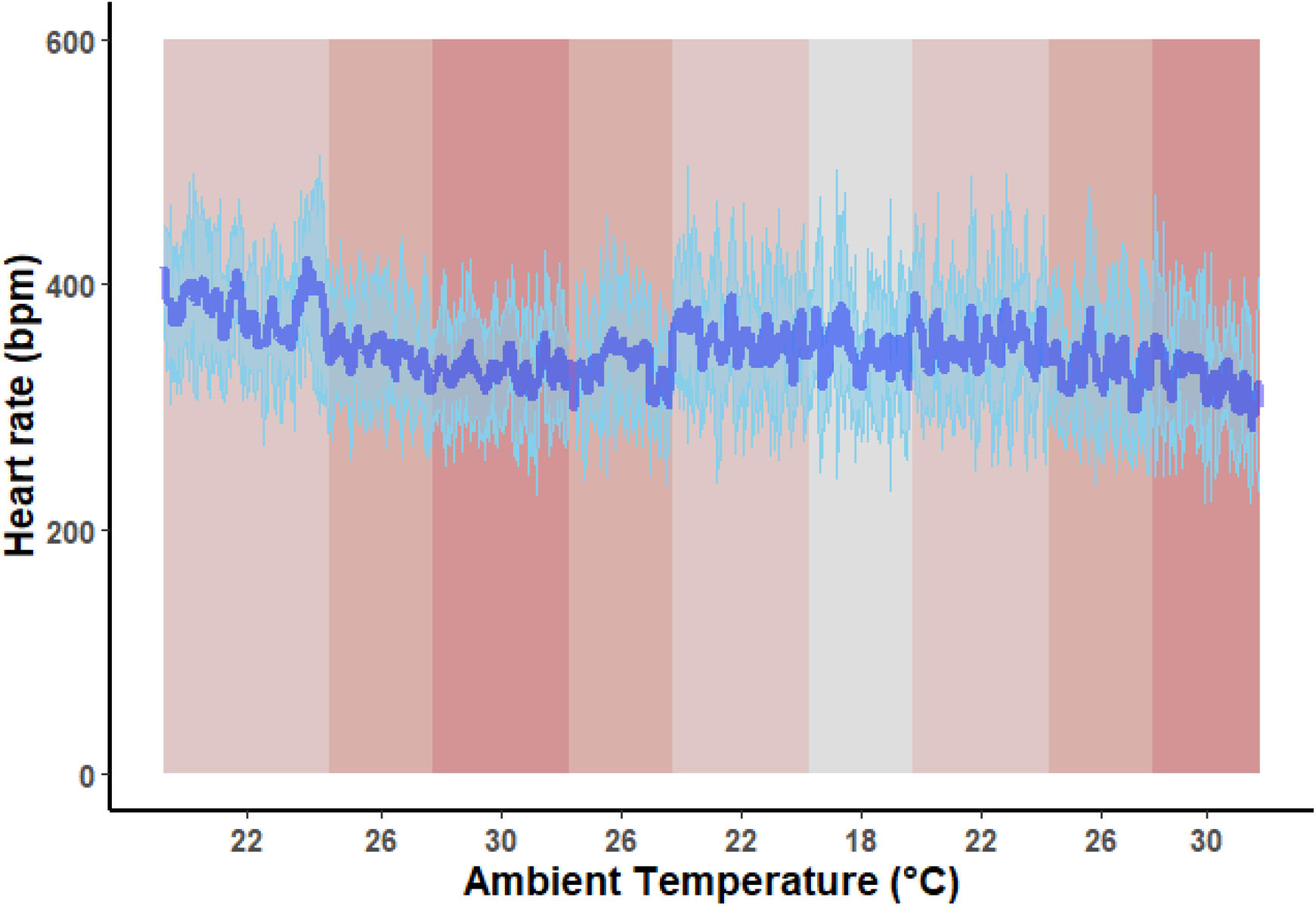
Heart rate was recorded hourly as ambient temperature varied and is shown here as a moving average over a 5-hour window. The moving average from 7 voles is shown as the dark blue line with the standard deviation of heart rate shown as the light blue ribbon behind it. Ambient temperature was varied continuously from 18° to 30° in 4° increments with 3-4 days spent at each stage. Ambient temperature is indicated on the x-axis and by shading, with darker colored panels indicating warmer ambient temperature.

### 3.2 Weekly Weights

In the high fat diet condition, at sexual maturity (PND45), CS animal body weights exceeded the average body weight of VD voles under the same diet condition. After PND45, the CS animal body weights dramatically increased and continued to increase in adulthood until sacrifice (PND75).

There were main effects of age (F(1,661.23) = 1125.89, p < 0.001) and sex (F(1,32.79) = 11.853, p = 0.002) on body weight. There were also interaction effects of birth mode X age (F(1,661.24) = 18.64, p < 0.001) and age X diet (F(1,66.23) = 12.23, p < 0.001), such that CS voles weighed more than VD counterparts as voles aged and the MC diet condition weighed more than the VC diet condition as voles aged (Table 1). Finally, there was also a trend toward a three-way interaction between age, birth mode, and diet, such that CS voles on the MC diet weighed the most as they aged (p = 0.056) (Figure 4).

**Table 1:** Type III Analysis of Variance Table with Satterthwaite’s Method for changes in body weight over time differentiated by birth mode and food condition. Significant Codes: 0 ‘**\*\*\***’ 0.001 ‘**\*\***’ 0.01 ‘**\***’ 0.05 ‘.’

|  | Sum Sq | Mean Sq | Num DF | DenDF | F value | Pr (>F) |  |
| --- | --- | --- | --- | --- | --- | --- | --- |
| Group | 34.9 | 34.9 | 1 | 91.10 | 1.5749 | 0.2127012 |  |
| Age | 24939.4 | 24939.4 | 1 | 661.23 | 1125.8964 | <2.2e-16 | *** |
| Sex | 262.6 | 262.6 | 1 | 32.79 | 11.8533 | 0.0015917 | ** |
| Diet | 9.7 | 9.7 | 1 | 91.40 | 0.4390 | 0.5092836 |  |
| Group: Age | 412.8 | 412.8 | 1 | 661.24 | 18.6364 | 1.824e-05 | *** |
| Group: Diet | 18.8 | 18.8 | 1 | 91.65 | 0.8486 | 0.3593797 |  |
| Age: Diet | 271.0 | 271.0 | 1 | 661.23 | 12.2323 | 0.0005011 | *** |
| Group:Age:Diet | 80.9 | 80.9 | 1 | 661.24 | 3.6515 | 0.0564525 | . |

#### 3.2.2 Length, Lee’s Index, and BMI at sacrifice

In terms of body length, there was a main effect of diet (F(1,26.09)=5.141, p = 0.032) such that voles in the MC diet condition were slightly longer. There were also trends towards greater length in males (p = 0.051) and CS voles (p = 0.065) (Figure 5). When we considered length as a covariate in our analysis of body weight, we observed a trend toward an interaction between birth mode and length (p = 0.064). Visual analysis of the scatterplot of length by weight revealed that CS voles had greater body weights, especially at relatively longer body lengths (Figure 5).

**Figure 5:**
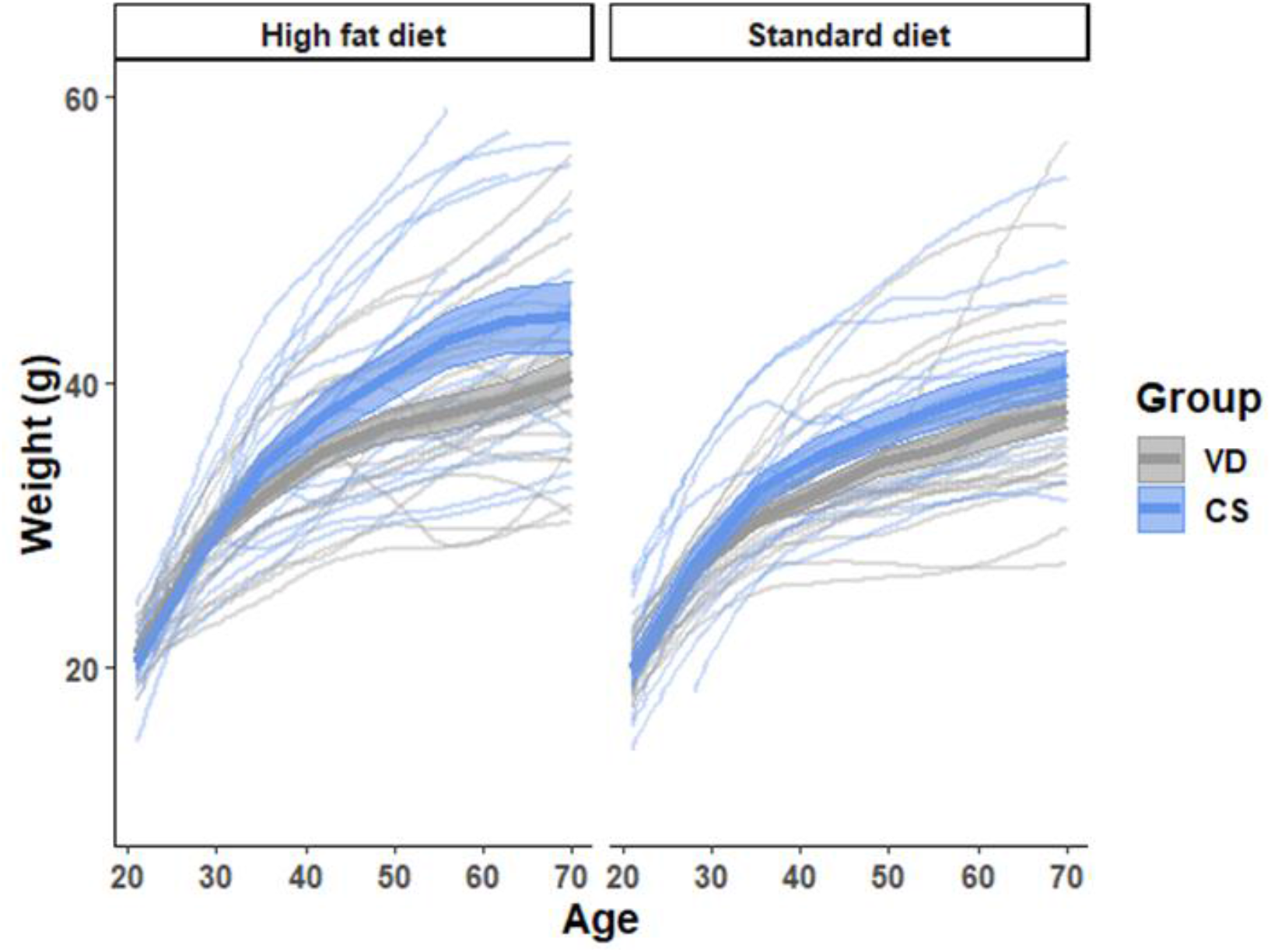
Changes in body weight of CS and VD voles of high-fat diet (left) and standard vole chow fed (right) prairie voles over time. VD: vaginal delivery; CS: cesarean delivery. Shaded regions depict 95% confidence intervals.

**Figure 6:**
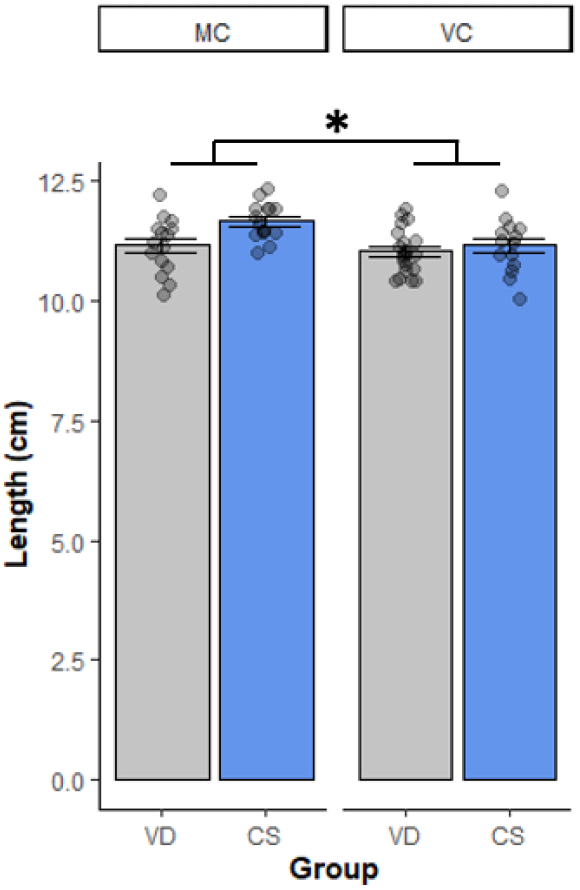
Length of vole at sacrifice differentiated by birth mode and diet condition.* denotes p<0.05. VD: vaginal delivery; CS: cesarean delivery; MC: mixed chow; VC: vole chow.

In terms of Lee’s index, we observed a main effect of sex (F(1,60) = 10.959, p = 0.002), such that males had greater scores than females (data not shown). In terms of BMI, we observed main effects of birth mode (F(1,60) = 5,7882, p = 0.019) and sex (F(1,60) = 20.934, p < 0.001) such that CS voles and males had greater BMI scores. However, post-hoc analyses did not reveal any significant differences in terms of birth mode between voles of the same sex (p > 0.05 for both comparisons; Figure 7).

**Figure 7:**
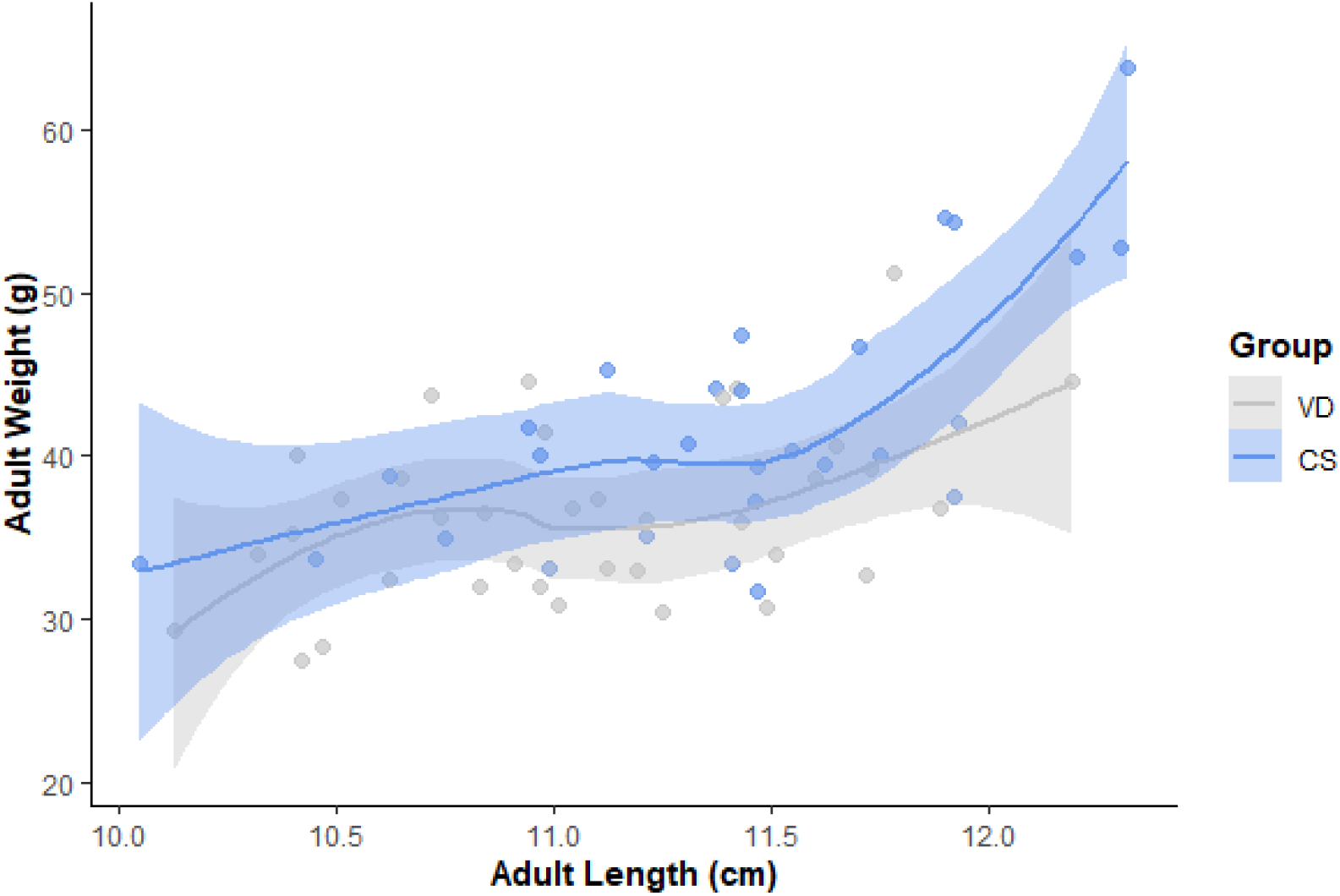
Length of vole at sacrifice as a function of weight. CS voles displayed greater body weights, especially at longer body lengths. Shaded regions depict 95% confidence intervals. Lee’s index and BMI were calculated as shown below:

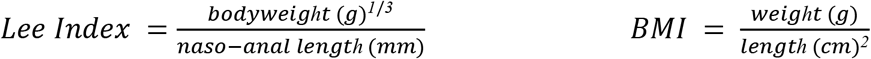

### 3.3 Adipose Tissue Weights

Both BAT and WAT fat pads were considered as proportional to the animal’s body weight.

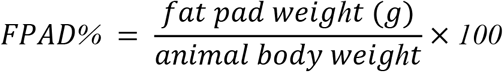

In terms of BAT, there were significant main effects of birth mode (F(1,28.474) = 5.18, p = 0.030) and diet (F(1,4.2, p = 0.049)), such that CS voles and VC voles had greater BAT as a percentage of body weight than counterparts. There was also a trend toward greater BAT weights in males than in females (p = 0.077)(data not shown). Post-hoc analyses revealed that within the VC diet condition, CS voles had greater percent BAT than VD counterparts (p = 0.036), whereas within the MC condition, CS/VD voles were not different, likely owing to small sample sizes in the MC+VD group (Figure 8).

**Figure 8:**
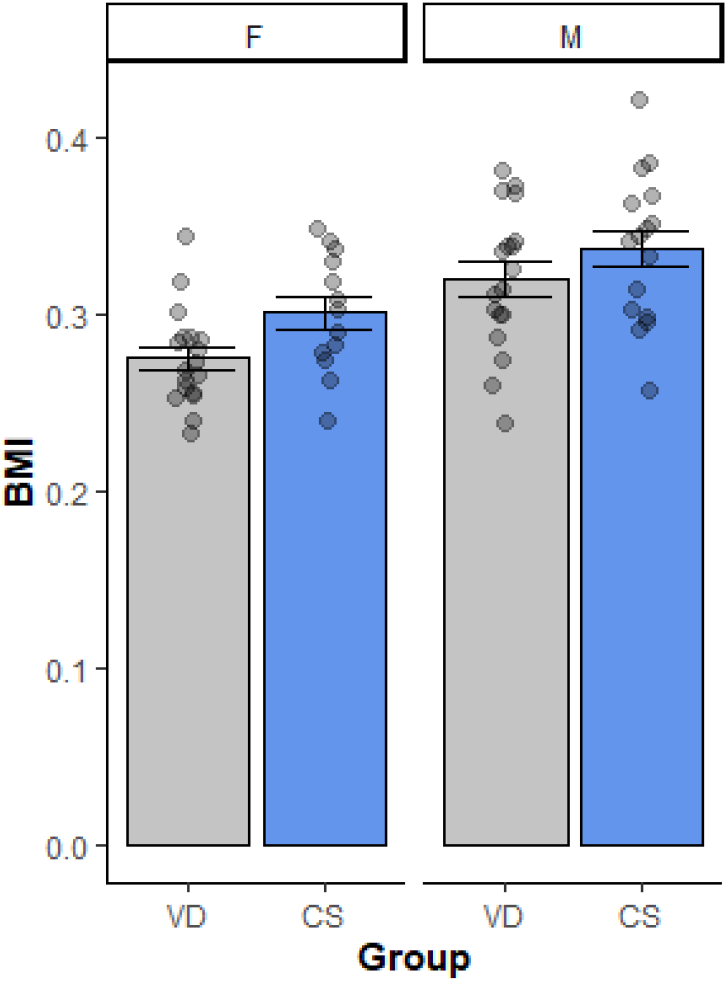
Body Mass Index (BMI) of male and female voles differentiated by birth mode at sacrifice. CS voles and males had greater BMI scores compared to their counterparts. F: female, M: male, VD: vaginal delivery, CS: cesarean delivery.

In terms of WAT, there was a main effect of diet (F(1,30.982) = 4.630, p = 0.039) such that animals in the VC diet condition had greater WAT as a percentage of body weight than counterparts (Figure 9).

**Figure 9:**
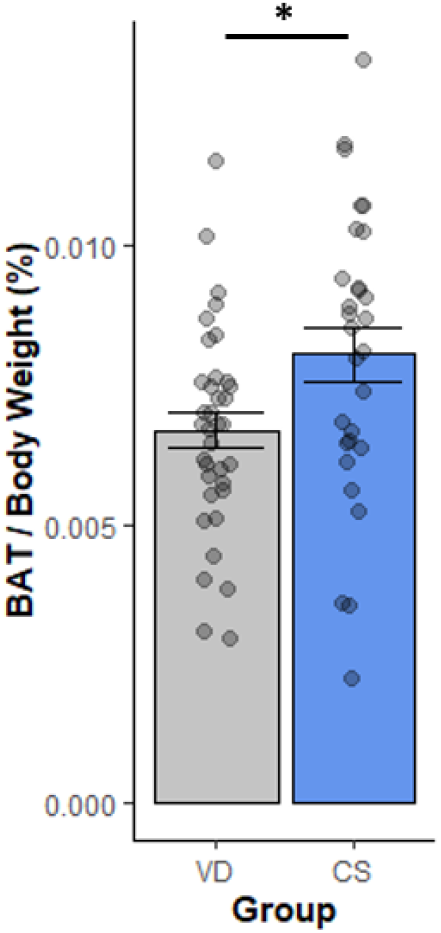
Bilateral BAT weights of animals at sacrifice (PND75), differentiated by birth mode. Data is collapsed across diet conditions and sex. (* denotes p < 0.05. VD: vaginal delivery; CS: cesarean delivery.)

### 3.4 Food Consumption

In terms of food consumption, there was a significant main effect of birth mode (F(1,13) = 25.44, p = 0.0149), such that CS voles consumed less food than VD counterparts (Figure 10). Because of small sample sizes, we excluded sex as a variable when analyzing food consumption. In terms of food consumption, we observed a main effect of birth mode (F(1,25.437) = 7.857, p = 0.015), such that CS cages consumed less food per animal than VD counterparts. Because of small sample sizes, we elected to not run post-hoc analyses.

**Figure 10:**
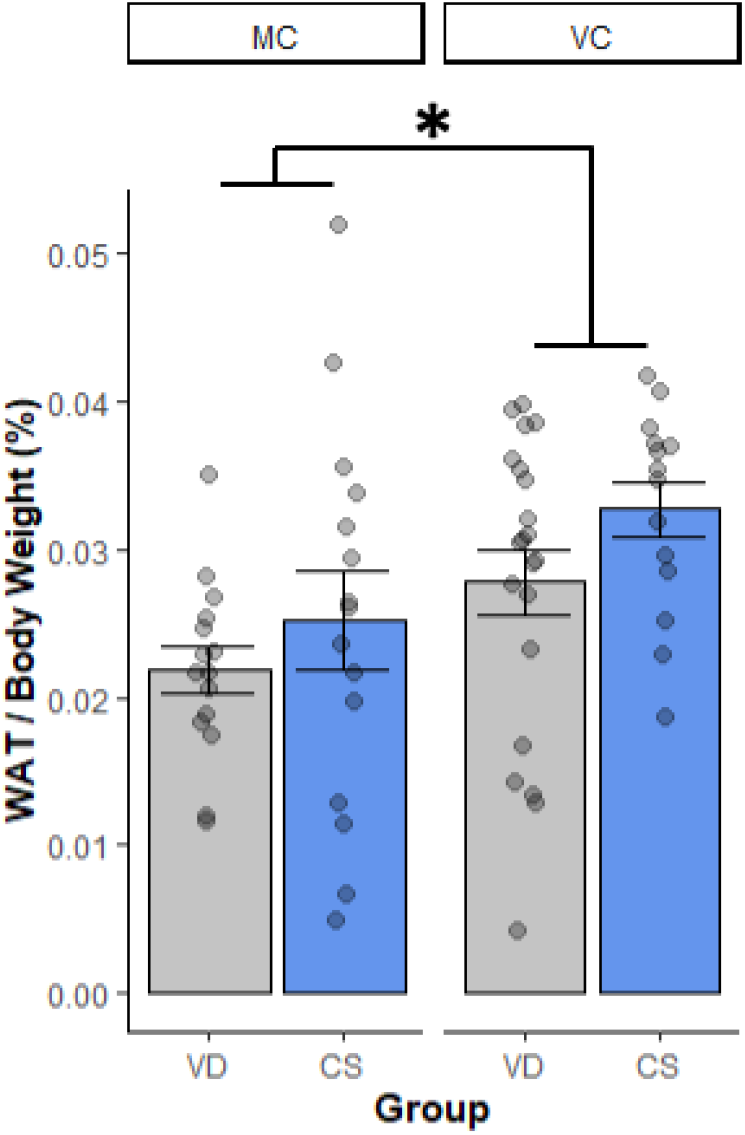
Bilateral WAT weights of animals at sacrifice, differentiated by birth mode and diet condition. Data is collapsed across sex. * denotes p<0.05. VD: vaginal delivery; CS: cesarean delivery.

### 3.5 Home Cage Activity

In terms of locomotor activity in the home cage, there was a main effect of birth mode (F(1,9.55), p = 0.025), such that CS voles showed greater locomotor activity than VD counterparts. Post-hoc analyses revealed significantly greater locomotor activity in CS voles compared to VD within the MC diet condition (p < 0.001). Due to small sample sizes, we did not consider post-hoc analysis for the VC diet condition (Figure 11).

**Figure 11:**
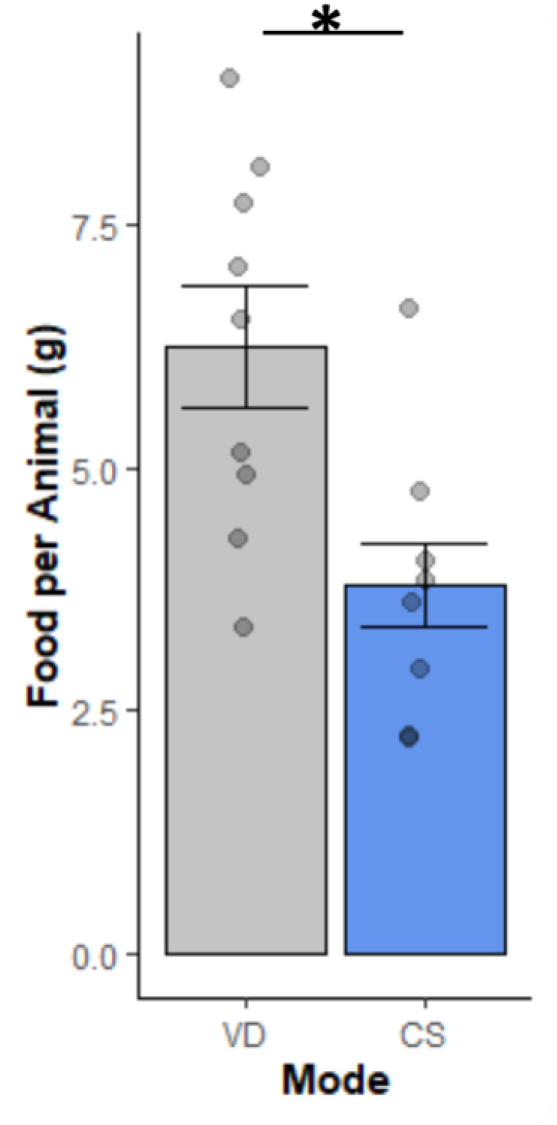
Food consumption per animal over a 24 hour period. Data shown is collapsed across diet condition groups. * denotes p<0.05. VD: vaginal delivery; CS: cesarean delivery.

**Figure 12:**
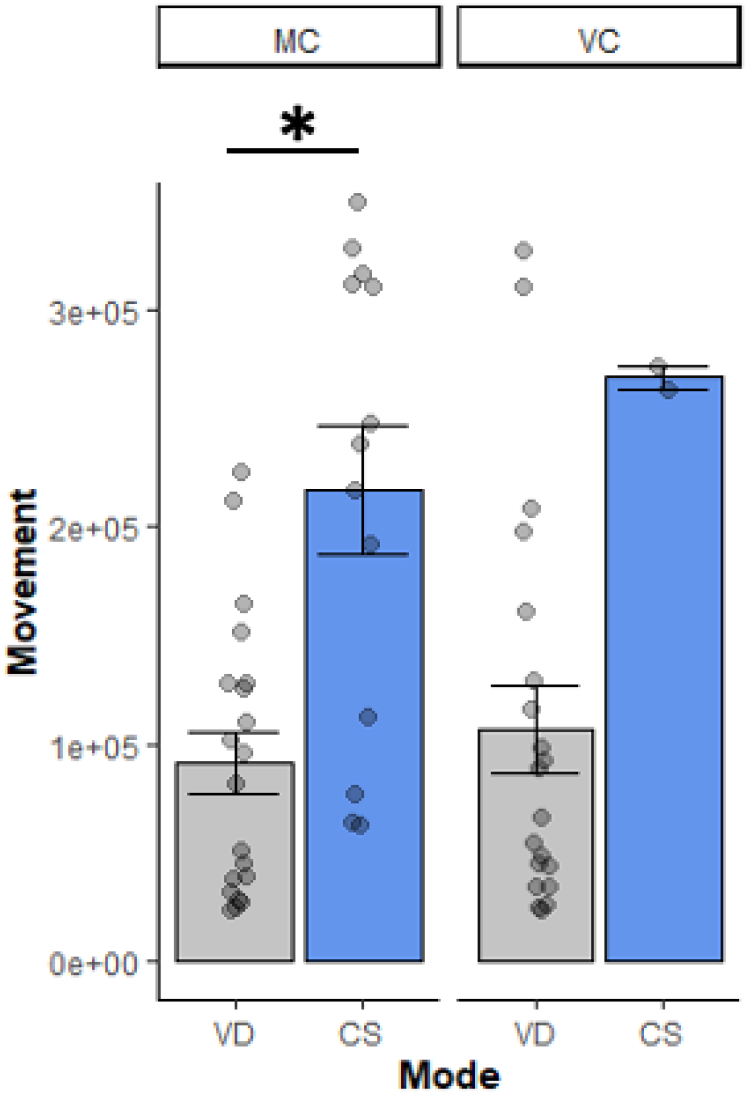
Repeated Measures ANOVA for Home cage activity at PND70 differentiated by birth mode and diet condition. * denotes p < 0.05. VD: vaginal delivery; CS: cesarean delivery; MC: mixed chow; VC: vole chow.

### 3.6 Sucrose Preference

Sucrose preference was determined by calculating the ratio of sucrose intake per total fluid intake.

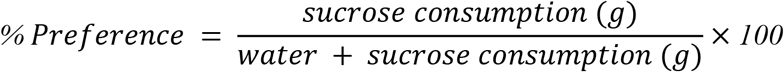

Several conditions, such as no drinking (when a litter does not drink either sucrose solution or water), over drinking (when the intake value of an animal is double that of the average for all voles in the study), or no determined preference (when an animal’s sucrose preference is <60%) were used as exclusion criteria for analysis (Liu et al., 2018). 5 cages from the PND30 dataset and 8 cages from the PND60 dataset were excluded from analysis due these criteria.

**Figure 13:**
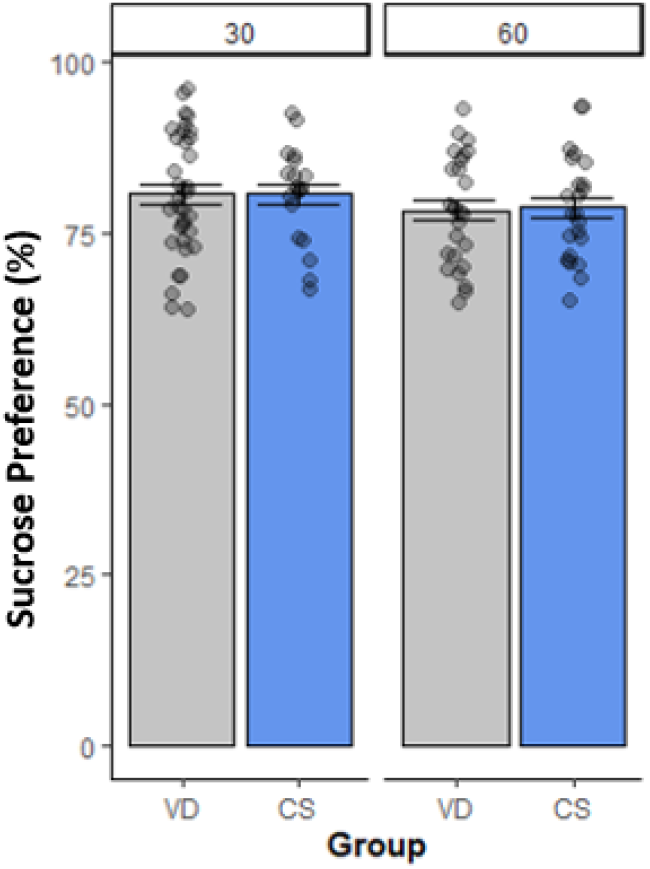
Repeated Measures ANOVA for sucrose preference at PND30 and PND70 differentiated by birth mode. Condensed over sex and diet condition. VD: vaginal delivery; CS: cesarean delivery.

We did not observe any significant difference between sucrose preference at PND30 and PND60 among diet conditions or birth mode groups. Only a single cage at a single time point showed <50% preference for sucrose, demonstrating that our voles overall prefered the sucrose solution over water.

## DISCUSSION

The results from this study established that 1) prairie voles tolerate a wide range of ambient temperatures and 2) that on metabolism after different birth modes and diet conditions found significant differences in body weight, food consumption, and activity. In experiment 1, we observed only a modest change in resting heart rate in response to changes in ambient temperature. Prairie vole heart rate declined 7-10% when ambient temperature was raised from 22° to 30°, changing from 363 to 327 bpm. During comparable changes in ambient temperature, lab mice show a dramatic shift in heart, changing from 603 to 363 bpm, a decline of 40% (Swoap et al., 2008). Thus, whereas mice find conventional housing conditions to be a chronic cold stress (Ganeshan & Chawla, 2017; Kokolus et al., 2013; Maloney et al., 2014a), the present findings support the view that prairie voles have a greater tolerance for variation in ambient temperature and thereby escape such chronic stress (Kenkel et al., 2021). Indeed, prior work has found that the thermoneutral range for prairie voles extends down into conventional ‘ room temperature’ (Beck and Anthony, 1971). Prairie voles have several adaptations suited to their year-round activity in their native habitat in the upper Great Plains of North America, including short tails and the genus’ eponymous small ears, both of which serve to reduce heat loss. Because the chronic cold stress posed by conventional housing conditions has led to spurious findings in studies of mice, these results suggest that the comparably sized prairie vole may offer advantages over traditional laboratory rodents (Kenkel et al., 2021). Species differences in thermoregulation are critically important to studies of metabolism, such as in the present experiment 2, because the chronic cold stress faced by mice and rats in conventional housing conditions acts as a major burden on the energy budget, ultimately slowing the development of weight gain.

Consistent with our predictions, CS offspring weighed more than their VD counterparts throughout development in both diet conditions (MC and VC). While gaining more weight across all groups, CS animals consumed less food over the single 24 hour testing period. CS animals also demonstrated greater home-cage locomotor activity than then their VD counterparts, most noticeably within the MC high-fat diet group during the single 2 hour testing period. All voles demonstrated a preference for the 1% sucrose solution, however there were no differences between groups. Post-mortem results indicate that CS animals had greater body weights especially at longer body lengths than VD animals. When calculating Body Mass Index (BMI), with the weights collected at sacrifice (PND75), CS animals in both diet groups had a larger score. The results from adipose tissue extractions used to reflect total adiposity were consistent with our predictions. In the VC diet condition, CS animals had a greater BAT as a percentage of body weight (%BAT) than VD counterparts. The WAT data as a percentage of body weight (%WAT) was inconsistent with our predictions, there was not a significant difference of %WAT among CS and VD groups. Among diet conditions, VC groups had a higher %WAT than the MC group. These findings demonstrate that birth mode plays an important role in the various anatomical and physiological systems associated with metabolism.

The collected pattern of results supports the hypothesis that birth via c-section leads to a metabolic change. The primary outcome of this study was that CS animals gained more weight than VD animals across both diet conditions and sexes; despite consuming less food and displaying increased locomotor activity. Thus, this difference in weight between groups cannot be attributed to inactivity or overeating and indicates alterations in metabolic processes. It is possible that this CS weight gain is due to increased muscle mass related to the observed increased locomotor activity, rather than the hypothesized adiposity. Furthermore the findings of increased %BAT in CS animals suggest a change in thermoregulation. Especially in small mammals, thermoregulation is a large component of energy expenditure which would have implications for metabolism. Very few studies of CS in humans have directly measured body composition beyond body weight. While unexpected, our results may still concur with the CS phenotype in humans.

This study also supports the notion that prairie voles avoid the chronic cold stress faced by traditional lab rodents in conventional housing conditions. We observed only a 7% change in heart rate among voles at 18° compared to 30°. Previous work examining the effect of ambient temperature on metabolism in mice found a >50% change in heart rate between 18° and 30°

(Swoap et al., 2008). This may relate to the fact that voles have thick coats of fur and are adapted to their native boreal habitats (Wunder, 1985; Wunder et al., 1977). Whereas thermoneutrality in mice is 30-32° (Cannon & Nedergaard, 2011; Hylander & Repasky, 2016), thermoneutrality in voles is ~25-30° (Beck & Anthony, 1971; Packard, 1968; Wunder et al., 1977). Vole thermoneutrality expands to 20-30° when nesting is considered (Beck & Anthony, 1971), however, nesting and group housing mice does little to reduce cold stress in mice (Maher et al., 2015). Ambient temperature is especially important in early life. Rearing rat pups at 22° leads to developmental programming such that food intake, thermoregulation, adiposity and sympathetic innervation all show enduring changes even after transfer to 28° housing (Young, 2006). Prairie voles’ biparental care is of particular relevance here, because fathers present a second source of warmth in the nest.

### 4.1.1 Possible Mechanisms

There are many physiological differences between vaginal delivery and elective cesarean section that could account for the results found in this study. There are birth hormones that, during labor, ‘surge’ through mother and fetus during vaginal delivery, including arginine vasopressin, estrogen, cortisol, progesterone, and oxytocin (Bridges,2015; Buckley,2015). Oxytocin is a reproductive hormone with many effects on the central nervous system and widespread activation of the parasympathetic nervous system. In a perinatal setting, maternal oxytocin increases before the onset of labor to increase the efficiency of the birth and induce contractions during labor.

Without the onset of labor, CS offspring are not exposed to the hormonal and inflammatory signals associated with birth hormones. A scheduled delivery via c-section reduces oxytocin release, including the maternal late-labor oxytocin surge and postpartum oxytocin peaks. The absence of oxytocin at birth could potentially lead to enduring effects on offspring development (Buckley, 2015; Prevost, 2014). Oxytocin has a widespread impact on adiposity, appetite, and thermoregulation throughout development. Oxytocin production has metabolic implications, such as reducing body weight and increasing glucose homeostasis (Ding et al., 2019).

Other studies have concluded that the gut microbiome is a possible mechanism impacting the metabolism of CS offspring. Birth via cesarean does not expose offspring to maternal microflora located in the vaginal canal. The absence of this exposure is believed to result in the reshaping of the offspring’s microflora makeup and diversity (Biasucci et al., 2008). Some studies have found that altered microflora composition has been linked with the development of obesity, however there are ongoing debates surrounding the factors underlying this relationship (Nieuwdorp, 2014). Further research is needed to identify whether the absence of hormonal surge or microflora exposure at birth is the critical mechanism contributing to the differences seen between CS and VD offspring in this study.

### 4.2 Limitations

There are several limitations to this research. We have observed the interactional effects of CS and high-fat diet supplementation on weight gain, but none of our current measures are complete assessments of obesity. We collected representative fat pad measures and body weight data to analyze body composition. In future studies, micro-CT scans will be used to better analyze percent adiposity of overall animal composition, especially visceral adiposity. This may explain the lack of findings surrounding measures of body composition through calculations like Lee’s Index or BMI. Next, home cage monitoring behavioral tests should be improved in future studies using passive monitors that do not impact nesting layout, food, and water access as well as allow us to record for longer periods of time.

The most severe limitation of this study was the measurement of caloric intake during representative food consumption periods. It is possible to make assumptions about caloric intake regarding the mixed chow using the caloric makeup of vole and mouse chow. However, we cannot discern the ratio of mouse to vole chow consumed within the mixed diet or high-fat diet condition, as food was given ad libitum and measured as a mixture. In future studies it will be necessary to measure how much of each component of the mixed chow is being consumed to better understand the exact nutritional profile of the high-fat diet.

#### 4.2.1 Future Research

Future research using a larger sample size within the home cage activity and food consumption is expected to further this exploration and provide valuable insights into the relationship between CS offspring and VD offspring in a high-fat diet condition. One essential addition to future studies of metabolism and birth mode under various diet conditions will be the assessment of animal preference. By understanding the levels of fiber, fat, and carbohydrates consumed, it will be possible to reach more informed conclusions regarding the impact of birth mode on dietary preferences.

Future studies will explore the neuroendocrine pathways surrounding birth, this can be done by targeting the idea of hormone surge at birth and the impact of hormones.To investigate oxytocin as the main mechanism in the metabolic changes discussed, it is possible to inject CS offspring with exogenous oxytocin at birth to mimic the hormonal surge administered during vaginal delivery and mitigate the metabolic outcomes due to CS. Conversely, a future study that tests the administration of an antagonist to VD offspring at birth may result in metabolic findings similar to CS offspring. With the addition of exogenous oxytocin and/or antagonist conditions, future studies may begin to more directly account for the hormonal changes during birth and record their effect on the metabolic factors explored in this paper (i.e. food composition, sucrose preference, body weights, adiposity, and home cage activity).

In these future studies, measures of home cage activity and locomotion will be augmented with repeated measures of home cage activity throughout adulthood, including longer periods of observation and decreased disruption to litters. The use of subcutaneous passive monitors (“PIT” tags) implanted at maturity may provide insight into thermoregulation of CS and VD litters under natural conditions. Another addition to make for future studies will be the inclusion of more advanced body composition measures. In particular, a micro-CT analysis of all animals at adulthood will assess body composition by measuring total adiposity, including subcutaneous and visceral fat content. With the addition of these more advanced data collection measures, we will work to understand the mechanisms behind the results collected in this study.

### 4.3 Conclusion

This experiment investigated metabolic changes associated with cesarean delivery. CS offspring were found to be more susceptible to increased weight gain than their VD counterparts, especially when both groups were given access to a high fat diet. This research lays the foundation on which to build an understanding of the metabolic implications of birth via cesarean section in an animal model. Future research will be done to elucidate the mechanisms involved in the relationship between metabolism and c-section. Given the widespread prevalence of c-sections and implication on metabolism, health and well-being.

## Notes

### Competing Interest Statement

The authors have declared no competing interest.

### Summary of Updates

This version includes experiment 1, where we tested the effects of ambient temperature on prairie vole metabolic rate. We found that prairie voles show a robust tolerance for variation in ambient temperature, with only a modest change in heart rate between 30 degrees and 22 degrees. This finding suggests that prairie voles avoid the chronic cold stress that impacts metabolic studies of traditional rodents housed in conventional conditions.

## REFERENCES

Almind K, Manieri M, Sivitz WI, Cinti S, Kahn CR. Ectopic brown adipose tissue in muscle provides a mechanism for differences in risk of metabolic syndrome in mice. Proc Natl Acad Sci U S A. 2007;104:2366–2371.

Bartoshuk LM, Duffy VB, Hayes JE, Moskowitz HR, Snyder DJ (2006) Psychophysics of sweet and fat perception in obesity: problems, solutions and new perspectives. Philos Trans R Soc Lond B Biol Sci 361: 1137–1148.

Beck, L. R., & Anthony, R. G. Metabolic and Behavioral Thermoregulation in the Long-Tailed Vole, Microtus longicaudus. Journal of Mammalogy, 52(2), 404–412 (1971).

Beura, L. K. et al. Normalizing the environment recapitulates adult human immune traits in laboratory mice. Nature 532, 512–516 (2016).

Bronwen Martin, Sunggoan Ji, Stuart Maudsley, and Mark P. Mattson “Control” laboratory rodents are metabolically morbid: Why it matters, March 1, 2010 doi:107(14)6127–6133 https://doi.org/10.1073/pnas.0912955107

Buckley, Sarah J. Hormonal Physiology of Childbearing: Evidence and Implications for Women, Babies, and Maternity Care. Washington, D.C.: Childbirth Connection Programs, National Partnership for Women & Families, January 2015.

Bridges et al., (2015) Front Neuroendocrinol. Front Neuroendocrinol. 2015 Jan; 0: 178–196. Published online 2014 Dec 10. doi: 10.1016/j.yfrne.2014.11.007

Cannon, B., & Nedergaard, J. (2004). Brown adipose tissue: function and physiological significance. Physiological reviews, 84(1), 277–359. https://doi.org/10.1152/physrev.00015.2003

Castillo-Ruiz, A., Mosley, M., Jacobs, A.J., Hoffiz, Y.C., Forger, N.G., 2018. Birth delivery mode alters perinatal cell death in the mouse brain. Proc.Natl.Acad.Sci.U.S.A. 115, 11826–11831. https://doi.org/10.1073/pnas.1811962115

Cui, X. et al. Thermoneutrality decreases thermogenic program and promotes adiposity in high-fat diet-fed mice. Physiological Reports 4, e12799 (2016).

C. Ding, M. K.-S. Leow, F. Magkos. Oxytocin in metabolic homeostasis: implications for obesity and diabetes management-Obesity-Diabetes Management/Etiology and Pathophysiology (2018) https://doi.org/10.1111/obr.12757

Eniola Sogunle, The association between cesarean section delivery and later life obesity in 21–24-year-olds in an Urban South African birth cohort PLoS One. 2019; 14(11): e0221379. Published online 2019 Nov 14. doi: 10.1371/journal.pone.0221379

Fadel et al., (2011) Postmortem and Perimortem Cesarean Section: Historical, Religious and Ethical Considerations J IMA. 2011 Dec; 43(3): 194–200. Published online 2012 Jan 23. doi: 10.5915/43-3-7099

Giacomo Biasucci, Belinda Benenati, Lorenzo Morelli, Elena Bessi, Günther Boehm, Cesarean Delivery May Affect the Early Biodiversity of Intestinal Bacteria, The Journal of Nutrition, Volume 138, Issue 9, September 2008, Pages 1796S–1800S, https://doi.org/10.1093/jn/138.9.1796S

Grinker J (1978) Obesity and sweet taste. Am J Clin Nutr 31: 1078–1087.

Hillebrand JJ, Langhans W, Geary N. Validation of computed tomographic estimates of intra-abdominal and subcutaneous adipose tissue in rats and mice. Obesity (Silver Spring) 18: 848 – 853, 2010. doi:10.1038/oby.2009.341

Hu, T., Mills, K. T., Yao, L., Demanelis, K., Eloustaz, M., Yancy, W. S., Jr, Kelly, T. N., He, J., & Bazzano, L. A. (2012). Effects of low-carbohydrate diets versus low-fat diets on metabolic risk factors: a meta-analysis of randomized controlled clinical trials. American journal of epidemiology, 176 Suppl 7(Suppl 7), S44–S54. https://doi.org/10.1093/aje/kws264

Keag O.E., Norman J., Stock S.J. Long-term risks and benefits associated with cesarean delivery for mother, baby, and subsequent pregnancies: Systematic review and meta-analysis. PLoS Med. 2018; 15: e1002494. Doi: 10.1371/journal.pmed.1002494

Kenkel, W. et al. Behavioral and epigenetic consequences of oxytocin treatment at birth. Science Advances 5, eaa v2244 (2019).

Kenkel, W. Birth signaling hormones and the developmental consequences of cesarean delivery. Journal of Neuroendocrinology n/a, e12912 (2020).

Kenkel, W., Gustison, M. L. & Beery, A. K. A Neuroscientist’s Guide to the Vole. Current Protocols 1, e175 (2021).

Kenkel, W. M., Paredes, J., Lewis, G. F., Yee, J. R., Pournajafi-Nazarloo, H., Grippo, A. J., Porges, S. W., & Carter, C. S.. Autonomic substrates of the response to pups in male prairie voles. PLoS One, 8(8), e69965. (2013).

Kenkel, W. M., Suboc, G., & Carter, C. S. Autonomic, behavioral and neuroendocrine correlates of paternal behavior in male prairie voles. Physiol Behav, 128, 252–259. (2014).

Kenkel, W. M., Yee, J. R., Porges, S. W., Ferris, C. F., & Carter, C. S. Cardioacceleration in alloparents in response to stimuli from prairie vole pups: The significance of thermoregulation. Behav Brain Res, 286, 71–79. (2015).

Kozhimannil, K. B., Hardeman, R. R., Attanasio, L. B., Blauer-Peterson, C., & O’Brien, M. (2013). Doula care, birth outcomes, and costs among Medicaid beneficiaries. American journal of public health, 103(4), e113–e121. https://doi.org/10.2105/AJPH.2012.301201

Kristy Townsend et al., Brown Fat Fuel Utilization and ThermogenesisTrends Endocrinol Metab. 2014 Apr; 25(4): 168–177. Published online 2014 Jan 2. doi: 10.1016/j.tem.2013.12.004

Lauren Jansen et al., 2013 First Do No Harm: Interventions During Childbirth J Perinat Educ. 2013 Spring; 22(2): 83–92. doi: 10.1891/1058-1243.22.2.83

Liu, X. et al. Brown Adipose Tissue Transplantation Reverses Obesity in Ob/Ob Mice. Endocrinology 156, 2461–2469 (2015).

Liu, MY., Yin, CY., Zhu, LJ. et al. Sucrose preference test for measurement of stress-induced anhedonia in mice. Nat Protoc 13, 1686–1698 (2018). https://doi.org/10.1038/s41596-018-0011-z

MacDorman FM, Mathews TJ, Declercq E. Trends in out-of-hospital births, 1990–2012. NCHS Data Brief No. 144 Hyattsville, MD: National Center for Health Statistics; 2014.

Maiken Scott, 2011 How did birth move from the home to the hospital, and back again? PBS NPR

Maloney, S. K., Fuller, A., Mitchell, D., Gordon, C. & Overton, J. M. Translating animal model research: Does it matter that our rodents are cold? Physiology (Bethesda, Md.) 29, 413–20 (2014).

Marian MacDorman et al., Trends and State Variations in Out-of-Hospital Births in the United States, 2004-2017 2011 Birth. 2019 Jun; 46(2): 279–288. Published online 2018 Dec 10. doi: 10.1111/birt.12411

Marie Prevost et al., 2014 Oxytocin in Pregnancy and the Postpartum: Relations to Labor and Its Management Front Public Health. 2014; 2: 1. Published online 2014 Jan 27. doi: 10.3389/fpubh.2014.00001

Masukume, G., Khashan, A. S., Morton, S., Baker, P. N., Kenny, L. C., & McCarthy, F. P. (2019). Cesarean section delivery and childhood obesity in a British longitudinal cohort study. PloS one, 14(10), e0223856. https://doi.org/10.1371/journal.pone.0223856

Montoya-Williams D., Lemas D.J., Spiryda L., Patel K., Carney O.O., Neu J., Carson T.L. The Neonatal Microbiome and Its Partial Role in Mediating the Association between Birth by Cesarean Section and Adverse Pediatric Outcomes. Neonatology. 2018; 114:103–111. doi: 10.1159/000487102

Nieuwdorp, M., Gilijamse, P. W., Pai, N., & Kaplan, L. M. (2014). Role of the microbiome in energy regulation and metabolism. Gastroenterology, 146(6), 1525–1533. https://doi.org/10.1053/j.gastro.2014.02.008

NIH Cesarean Section-A Brief History U.S. National Library of Medicine https://www.nlm.nih.gov/exhibition/cesarean/part1.html

Novelli, E. L., Diniz, Y. S., Galhardi, C. M., Ebaid, G. M., Rodrigues, H. G., Mani, F., Fernandes, A. A., Cicogna, A. C., & Novelli Filho, J. L. (2007). Anthropometrical parameters and markers of obesity in rats. Laboratory animals, 41(1), 111–119. https://doi.org/10.1258/002367707779399518

Potretzke et al., The Prairie Vole Model of Pair-Bonding and Its Sensitivity to Addictive SubstancesFront. Psychol., 06 November 2019 | https://doi.org/10.3389/fpsyg.2019.02477

Rogers, P., & Webb, G. P. (1980). Estimation of body fat in normal and obese mice. The British journal of nutrition, 43(1), 83–86. https://doi.org/10.1079/bjn19800066

Stemmer, K. et al. Thermoneutral housing is a critical factor for immune function and diet-induced obesity in C57BL/6 nude mice. Int J Obes (Lond) 39, 791–797 (2015).

Sbierski-Kind, J. et al. Distinct Housing Conditions Reveal a Major Impact of Adaptive Immunity on the Course of Obesity-Induced Type 2 Diabetes. Front. Immunol. 9, (2018).

Soos S, Balasko M, Jech-Mihalffy A, Szekely M, Petervari E. Anorexic vs. metabolic effects of central leptin infusion in rats of various ages and nutritional states. J Mol Neurosci 41: 97–104, 2010. Doi: 10.1007/s12031-009-9294-4.

Swoap, S. J., Li, C., Wess, J., Parsons, A. D., Williams, T. D., & Overton, J. M. Vagal tone dominates autonomic control of mouse heart rate at thermoneutrality. American Journal of Physiology-Heart and Circulatory Physiology, 294(4), H1581–H1588 (2008).

Tang H, Vasselli JR, Wu EX, Boozer CN, Gallagher D. High-resolution magnetic resonance imaging tracks changes in organ and tissue mass in obese and aging rats. Am J Physiol Regul Integr Comp Physiol 282: R890–R899, 2002. doi:10.1152/ajpregu.0527.2001.

World Health Organization Human Reproduction Programme, 10 April 2015 (2015). WHO Statement on cesarean section rates. Reproductive health matters, 23(45), 149–150. https://doi.org/10.1016/j.rhm.2015.07.007

World Health Organization. Obesity and Overweight. Fact sheet N° 311. (Online). http://www.who.int/mediacentre/factsheets/fs311/en/ [12 Dec. 2017].

Yuan, C., Gaskins, A. J. & Blaine, A. I. Association between cesarean birth and risk of obesity in offspring in childhood, adolescence, and early adulthood. JAMA Pediatrics 170, e162385 (2016).

Zachariassen, L. F. et al. Cesarean Section Induces Microbiota-Regulated Immune Disturbances in C57BL/6 Mice. J Immunol 202, 142–150 (2018).

Beck, L. R., & Anthony, R. G. (1971). Metabolic and Behavioral Thermoregulation in the Long-Tailed Vole, Microtus longicaudus. Journal of Mammalogy, 52(2), 404–412. https://doi.org/10.2307/1378682

Cannon, B., & Nedergaard, J. (2011). Nonshivering thermogenesis and its adequate measurement in metabolic studies. The Journal of Experimental Biology, 214(Pt 2), 242–253. https://doi.org/10.1242/jeb.050989

Corrigan, J. K., Ramachandran, D., He, Y., Palmer, C. J., Jurczak, M. J., Chen, R., Li, B., Friedline, R. H., Kim, J. K., Ramsey, J. J., Lantier, L., McGuinness, O. P., & Banks, A. S. (2020). A big-data approach to understanding metabolic rate and response to obesity in laboratory mice. ELife, 9(Journal Article), e53560. https://doi.org/10.7554/eLife.53560

Cui, X., Nguyen, N. L. T., Zarebidaki, E., Cao, Q., Li, F., Zha, L., Bartness, T., Shi, H., & Xue, B. (2016). Thermoneutrality decreases thermogenic program and promotes adiposity in high-fat diet-fed mice. Physiological Reports, 4(10), e12799. https://doi.org/10.14814/phy2.12799

Feldmann, H. M., Golozoubova, V., Cannon, B., & Nedergaard, J. (2009). UCP1 ablation induces obesity and abolishes diet-induced thermogenesis in mice exempt from thermal stress by living at thermoneutrality. Cell Metabolism, 9(2), 203–209. https://doi.org/10.1016/j.cmet.2008.12.014

Ganeshan, K., & Chawla, A. (2017). Warming the mouse to model human diseases. Nature Reviews Endocrinology, 13(8), 458–465. https://doi.org/10.1038/nrendo.2017.48

Giles, D. A., Moreno-Fernandez, M. E., Stankiewicz, T. E., Graspeuntner, S., Cappelletti, M., Wu, D., Mukherjee, R., Chan, C. C., Lawson, M. J., Klarquist, J., Sünderhauf, A., Softic, S., Kahn, C. R., Stemmer, K., Iwakura, Y., Aronow, B. J., Karns, R., Steinbrecher, K. A., Karp, C. L.,… Divanovic, S. (2017). Thermoneutral housing exacerbates nonalcoholic fatty liver disease in mice and allows for sex-independent disease modeling. Nature Medicine, 23(7), 829–838. https://doi.org/10.1038/nm.4346

Hankenson, F. C., Marx, J. O., Gordon, C. J., & David, J. M. (2018). Effects of Rodent Thermoregulation on Animal Models in the Research Environment. Comparative Medicine, 68(6), 425–438. https://doi.org/10.30802/AALAS-CM-18-000049

Hylander, B. L., & Repasky, E. A. (2016). Thermoneutrality, Mice, and Cancer: A Heated Opinion. Trends in Cancer, 2(4), 166–175. https://doi.org/10.1016/j.trecan.2016.03.005

Kokolus, K. M., Capitano, M. L., Lee, C.-T., Eng, J. W.-L., Waight, J. D., Hylander, B. L., Sexton, S., Hong, C.-C., Gordon, C. J., Abrams, S. I., & Repasky, E. A. (2013). Baseline tumor growth and immune control in laboratory mice are significantly influenced by subthermoneutral housing temperature. Proceedings of the National Academy of Sciences, 110(50), 20176–20181. https://doi.org/10.1073/pnas.1304291110

Maher, R. L., Barbash, S. M., Lynch, D. V., & Swoap, S. J. (2015). Group housing and nest building only slightly ameliorate the cold stress of typical housing in female C57BL/6J mice. American Journal of Physiology-Regulatory, Integrative and Comparative Physiology, 308(12), R1070–R1079. https://doi.org/10.1152/ajpregu.00407.2014

Maloney, S. K., Fuller, A., Mitchell, D., Gordon, C., & Overton, J. M. (2014a). Translating Animal Model Research: Does It Matter That Our Rodents Are Cold? Physiology, 29(6), 413–420. https://doi.org/10.1152/physiol.00029.2014

Maloney, S. K., Fuller, A., Mitchell, D., Gordon, C., & Overton, J. M. (2014b). Translating animal model research: Does it matter that our rodents are cold? Physiology (Bethesda), 29(6), 413–420. https://doi.org/10.1152/physiol.00029.2014

Packard, G. C. (1968). Oxygen Consumption of Microtus Montanus in Relation to Ambient Temperature. Journal of Mammalogy, 49(2), 215–220. https://doi.org/10.2307/1377977

Stemmer, K., Kotzbeck, P., Zani, F., Bauer, M., Neff, C., Müller, T. D., Pfluger, P. T., Seeley, R. J., & Divanovic, S. (2015). Thermoneutral housing is a critical factor for immune function and diet-induced obesity in C57BL/6 nude mice. International Journal of Obesity (2005), 39(5), 791–797. https://doi.org/10.1038/ijo.2014.187

Swoap, S. J., Li, C., Wess, J., Parsons, A. D., Williams, T. D., & Overton, J. M. (2008). Vagal tone dominates autonomic control of mouse heart rate at thermoneutrality. American Journal of Physiology-Heart and Circulatory Physiology, 294(4), H1581–H1588. https://doi.org/10.1152/ajpheart.01000.2007

Wunder, B. A. (1985). Energetics and thermoregulation. In Biology of new world Microtus (Vol. 8, pp. 812–844). Am. Soc. Mammalogist Lawrence, Kansas.

Wunder, B. A., Dobkin, D. S., & Gettinger, R. D. (1977). Shifts of thermogenesis in the prairie vole (Microtus ochrogaster). Oecologia, 29(1), 11–26. https://doi.org/10.1007/BF00345359

Young, J. B. (2006). Developmental origins of obesity: A sympathoadrenal perspective. International Journal of Obesity, 30(S4), S41–S49. https://doi.org/10.1038/sj.ijo.0803518

